# Glutamine supports the protection of tissue cells against the damage caused by cholesterol-dependent cytolysins from pathogenic bacteria

**DOI:** 10.1101/679068

**Authors:** Matthew L. Turner, Sian E. Owens, I. Martin Sheldon

**Affiliations:** Institute of Life Science, Swansea University Medical School, Swansea University, Swansea, United Kingdom

## Abstract

Pathogenic bacteria often damage tissues by secreting toxins that form pores in cell membranes, and the most common pore-forming toxins are cholesterol-dependent cytolysins. During bacterial infections, glutamine becomes a conditionally essential amino acid, and glutamine is an important nutrient for immune cells. However, the role of glutamine in protecting tissue cells against pore-forming toxins is unclear. Here we tested the hypothesis that glutamine supports the protection of tissue cells against the damage caused by cholesterol-dependent cytolysins. Stromal and epithelial cells were sensitive to damage by cholesterol-dependent cytolysins, pyolysin and streptolysin O, as determined by leakage of potassium and lactate dehydrogenase from cells, and reduced cell viability. However, glutamine helped protect cells against cholesterol-dependent cytolysins because glutamine deprivation increased the leakage of lactate dehydrogenase and reduced the viability of cells challenged with cytolysins. Without glutamine, stromal cells challenged with pyolysin leaked lactate dehydrogenase (control vs. pyolysin, 2.6 ± 0.6 vs. 34.4 ± 4.5 AU, n = 12), which was more than three-fold the leakage from cells supplied with 2 mM glutamine (control vs. pyolysin, 2.2 ± 0.3 vs. 9.4 ± 1.0 AU). The cytoprotective effect of glutamine was not dependent on glutaminolysis, replenishing the Krebs cycle via succinate, changes in cellular cholesterol, or regulators of cell metabolism (AMPK and mTOR). In conclusion, although the mechanism remains elusive, we found that glutamine supports the protection of tissue cells against the damage caused by cholesterol-dependent cytolysins from pathogenic bacteria.

## Introduction

Animals defend themselves against bacterial infections using the complimentary strategies of resistance and tolerance [1–3]. Resistance is the ability to limit the pathogen burden, usually by employing the immune system to kill bacteria. Tolerance is the ability to limit the severity of disease caused by a given pathogen burden, usually by limiting the damage caused by bacteria. Bacteria often damage tissue cells by secreting toxins that form pores in the cell membrane, and the most common pore-forming toxins are cholesterol-dependent cytolysins [4–7]. During bacterial infections, the cells of the immune system use glutamine as a key nutrient to support inflammatory responses [8–10]. However, the role of glutamine in protecting tissue cells against the damage caused by cholesterol-dependent cytolysins is unclear.

Cholesterol-dependent cytolysins include pyolysin secreted by *Trueperella pyogenes*, which causes purulent infections in cattle and swine, such as postpartum uterine disease and abscesses, and streptolysin O (SLO) secreted by beta-hemolytic group A *Streptococci*, which causes pharyngitis and impetigo in children [11–14]. These cytolysins bind cholesterol-rich areas in tissue cell membranes, where they form 30 nm diameter pores. The membrane pores lead to leakage of potassium ions from cells within minutes, and further cell damage is evidenced by leakage of proteins, such as lactate dehydrogenase (LDH), from the cytoplasm and ultimately cell death [6, 15]. Tissue cells counter the damage by activating stress responses and transitioning to a quiescent metabolic state [5, 6]. The effect of metabolism on cytoprotection against cholesterol-dependent cytolysins is largely unexplored. However, an intriguing observation is that the metabolic stress of lactation in dairy cattle increases the risk of postpartum uterine disease associated with *T. pyogenes* [16–19], probably by impairing the ability of the endometrial tissue to tolerate the presence of bacteria [20]. We therefore proposed that the availability of nutrients might affect the ability of tissue cells to protect themselves against cholesterol-dependent cytolysins.

Cells use glucose and glutamine to supply most of their energy [21–23]. Glycolysis converts glucose to pyruvate to feed the Krebs cycle, whilst glutaminase converts glutamine to glutamate to replenish the Krebs cycle [9, 24]. Glutamine is an abundant non-essential amino acid, with about 0.7 mM glutamine in human peripheral plasma and 0.25 mM in bovine plasma [8, 25]. However, glutamine becomes a conditionally essential amino acid after injury or infection, and glutamine fosters immune cell inflammatory responses [8, 9, 26, 27]. As glutamine is a key nutrient, our aim was to test the hypothesis that glutamine supports the protection of tissue cells against the damage caused by cholesterol-dependent cytolysins.

## Results

### Pyolysin damages stromal cells

We used primary bovine endometrial stromal cells and pyolysin to study cytoprotection because these tissue cells are the principal target for pyolysin [14]; and, unlike other cholesterol-dependent cytolysins, pyolysin does not require thiol-activation [13]. Pyolysin formed pores in the stromal cells, as determined by the loss of intracellular potassium within 5 min (Fig 1A). Furthermore, a 2 h challenge with pyolysin damaged the stromal cells, as determined by reduced cell viability (Fig 1B) and leakage of lactate dehydrogenase (LDH) from the cytosol into cell supernatants (Fig 1C). We chose a 2 h pyolysin challenge based on previous kinetic studies where 50% of endometrial stromal cells were perforated after 2 h [14]. Furthermore, the 2 h challenge reduces the likelihood of confounding cell protection with immune responses to cytolysins, which are usually evident after 2 h in immune cells [28].

**Fig 1.**
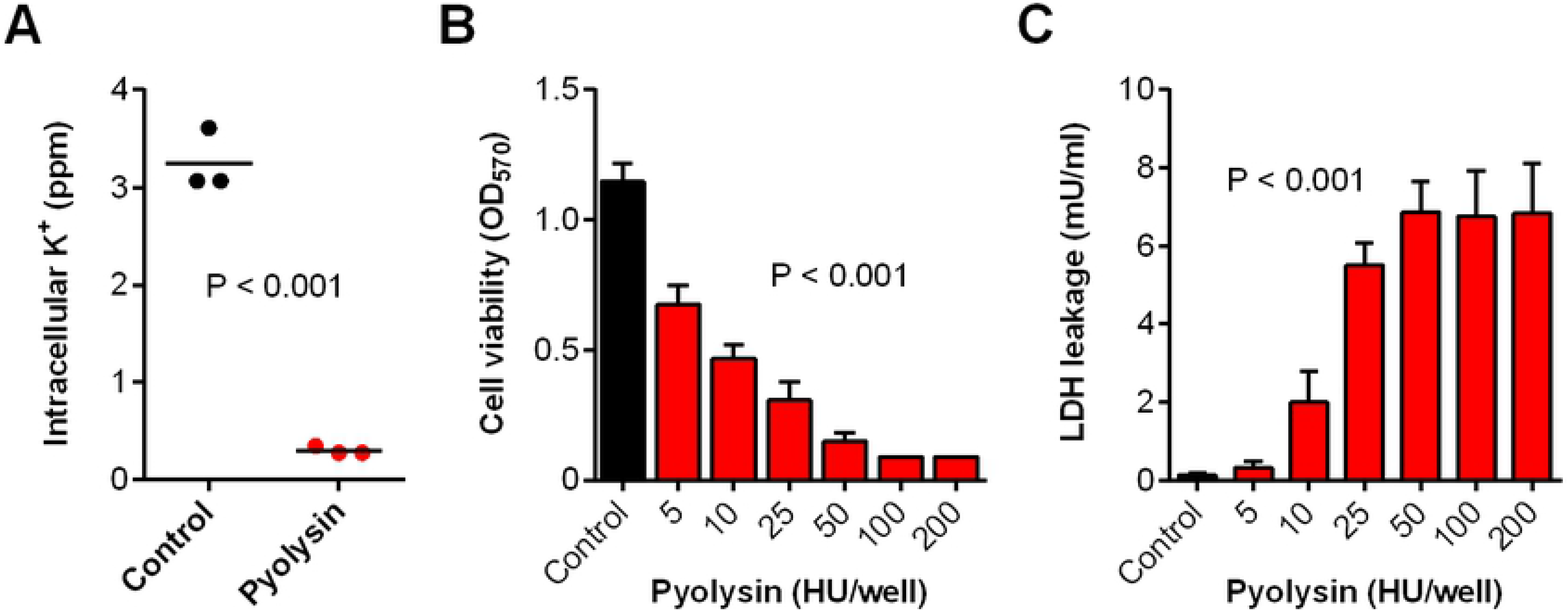
Cytolytic activity of pyolysin. (A) Bovine endometrial stromal cells were challenged for 5 min with control serum-free medium (•) or medium containing pyolysin 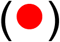, and potassium was measured in cell lysates. Data are presented using cells from 3 animals and the horizontal line represents the mean; data were analyzed by t-test. (B, C) Stromal cells were challenged for 2 h with control serum-free medium (▀) or medium with the indicated concentrations of pyolysin 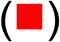; cell viability was determined by MTT assay (B) and LDH leakage evaluated by measuring LDH in the cell supernatants (C). Data are presented as mean (SEM) using cells from 4 animals; data were analyzed by ANOVA and P values are reported.

### Glutamine supports stromal cell protection against pyolysin

Cells are usually cultured in media containing 2 mM glutamine, which is eight fold higher than the plasma concentration of glutamine in cows [25]. To examine if the availability of glutamine affected cytoprotection against pyolysin, we cultured stromal cells for 24 h in serum-free media containing an excess of glucose (11.1 mM) with a range of concentrations of glutamine (0 to 2 mM), and then challenged the cells for 2 h with control medium or 10 HU/well pyolysin. We used serum-free medium because cholesterol in serum can bind cholesterol-dependent cytolysins, and to limit glutamine-dependent differences in cell growth. Irrespective of glutamine availability, pyolysin caused pore formation, as determined by loss of intracellular potassium within 5 min (Fig 2A; two-way ANOVA, n = 3 animals, P < 0.001). We next evaluated the effect of glutamine on pyolysin-induced cell damage by determining the leakage of LDH into cell supernatants. We did not use the mitochondrial-dependent MTT assay for cell viability here because differences in glutamine availability affect cell growth and mitochondrial function [8]. Furthermore, to account for differences in cell growth, we measured cellular DNA at the end of each experiment and normalized the leakage of LDH into cell supernatants using the control-challenge cells. Limiting the availability of glutamine increased the accumulation of LDH in supernatants when cells were challenged with pyolysin (Fig 2B; two-way ANOVA; n = 4 animals, P < 0.001). The increased leakage of LDH in stromal cells cultured without glutamine, compared with 2 mM glutamine, was evident from 15 min after pyolysin challenge (Fig 2C). When cells were examined by light microscopy, cells cultured with 2 mM glutamine and challenged with pyolysin showed some damage but usually maintained defined cell boundaries, whereas most cells were misshapen if they were deprived of glutamine and challenged with pyolysin (Fig 2D). Staining actin with phalloidin also showed that when challenged with pyolysin the cytoskeletal was more disrupted in cells deprived of glutamine than cells cultured with glutamine (Fig 2E). Finally, as primary cells often vary in their biological response, we verified our observations using stromal cells collected from 12 independent animals; cells cultured without glutamine leaked more than three times the LDH from cells supplied with 2 mM glutamine (Fig 2F).

**Fig 2.**
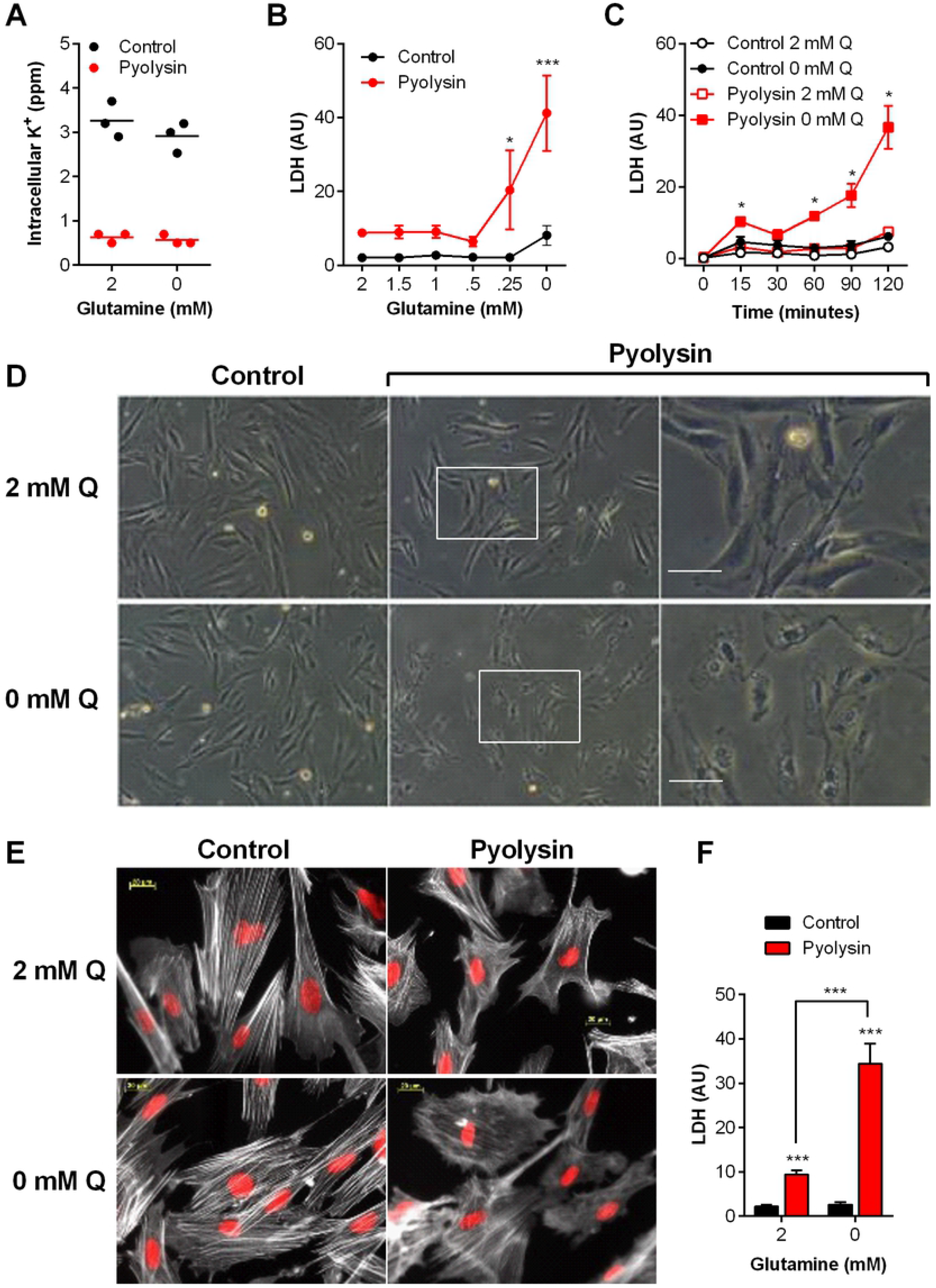
Glutamine is cytoprotective against pyolysin. (A) Bovine endometrial stromal cells were cultured for 24 h in medium containing 2 mM glutamine (2) or without glutamine (0), and challenged for 5 min with control medium (•) or pyolysin 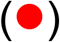. Intracellular potassium was determined by flame photometry. Data are from 3 animals, with a horizontal line indicating the mean. (B) Cells were cultured in medium containing the indicated concentrations of glutamine for 24 h, and challenged for 2 h with control medium (•) or 10 HU pyolysin 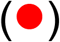, and LDH measured in supernatants and normalized to cellular DNA in the control challenge. Data are mean (SEM) of 4 animals, analyzed by ANOVA with Dunnett’s post hoc test; values differ from 2 mM glutamine pyolysin challenge, *** P <0.001, * P < 0.05. (C) Stromal cells were cultured with 2 mM glutamine (Q, open symbols) or without glutamine (0 mM Q, filled symbols) for 24 h, and challenged for the indicated times with control medium or pyolysin; LDH leakage was measured in supernatants and normalized to cellular DNA in the control challenge. Data are mean (SEM) of 4 animals; analyzed by ANOVA with Dunnett’s post hoc test; values differ from 2 mM glutamine pyolysin challenge, * P < 0.05. (D) Stromal cells were cultured with glutamine (2 mM Q) or without glutamine (0 mM Q) for 24 h, and challenged for 2 h with control medium or pyolysin. Transmitted light micrographs of cells were captured at the end of the experiment; right column represents magnification of boxed areas from middle column; scale bar 10 μm; images are representative of cells from 4 animals. (E) Cells were also stained with phalloidin to visualize F-actin (white) and fluorescent microscope images collected; nuclei are red; scale bars are 20 μm; images are representative of cells from 4 animals. (F) Cells were cultured for 24 h in medium containing 2 mM glutamine (2) or without glutamine (0), and challenged for 2 h with control medium (▀) or 10 HU pyolysin 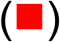, and LDH measured in supernatants and normalized to cellular DNA in the control challenge. Data are mean (SEM) of 12 animals, analyzed by ANOVA with Bonferroni post hoc test; values differ from 2 mM glutamine pyolysin challenge, *** P <0.001.

We examined the possibility that glutamine deprivation might increase LDH leakage because there was more intracellular LDH if glutamine-deprived cells used lactate as an alternative metabolic substrate to glutamine. However, LDH activity in cell lysates was similar after 24 h culture of cells with or without glutamine (Fig 3A; independent t-test, P = 0.65, n = 4 animals). We also considered the possibility that glutamine might bind to pyolysin or neutralize the activity of pyolysin. To test this possibility we exploited horse red blood cells, which are highly sensitive to hemolysis caused by cholesterol-dependent cytolysins [14]. However, incubating glutamine with pyolysin did not affect hemolysis, whereas incubating pyolysin with cholesterol, as a control, reduced hemolysis, as expected for a cholesterol-dependent cytolysin (Fig 3B).

**Fig 3.**
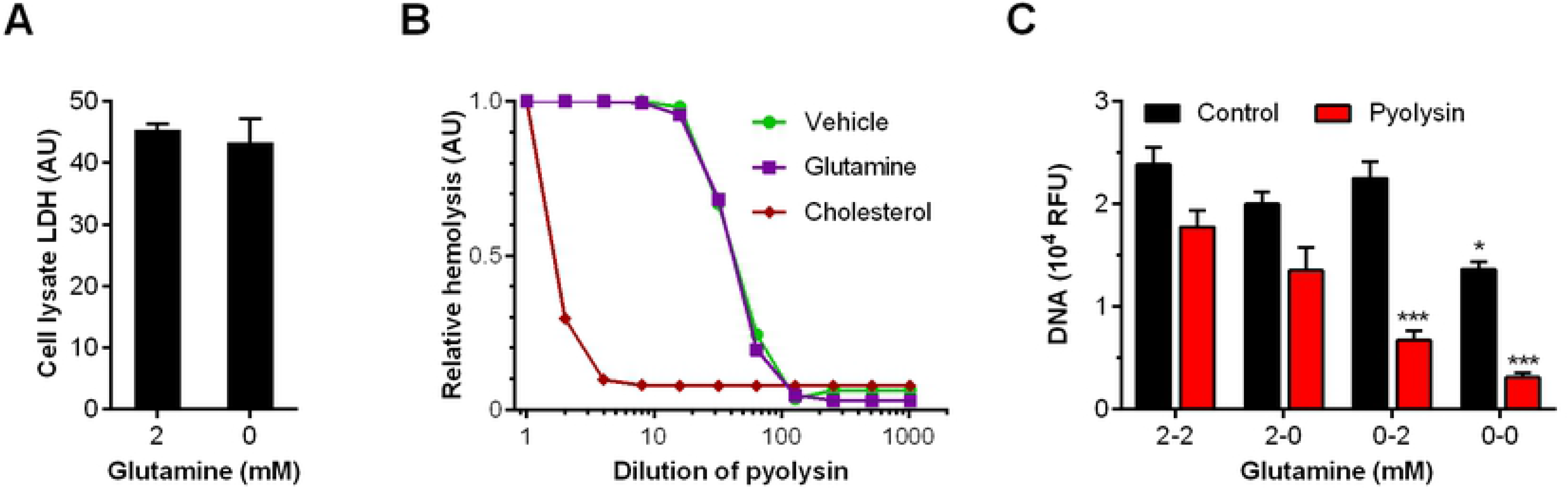
Glutamine is cytoprotective but does not alter intracellular LDH or bind pyolysin. (A) Whole cell lysates were collected after 24 h treatment with 2 mM (2) or no glutamine (0), and intracellular LDH abundance measured, and normalized to cellular DNA. (B) Cytolysis of red blood cells, as determined by hemolysis assay, when treated with a serial dilution of pyolysin, after prior incubation of the pyolysin for 1 h with vehicle 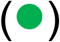, 2 mM glutamine 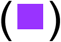, or 1 μM cholesterol 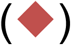 used as a positive control to bind to pyolysin. Data are presented as mean of 2 experiments, with 2 replicates per treatment. (C) Cells were cultured in serum-free medium containing 2 mM (2) or without glutamine (0) for 24 h before challenge with control medium (▀) or pyolysin 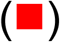 for 2 h. The media were then replenished with medium containing 2 mM glutamine (2) or without glutamine (0) for a further 24 h and cellular DNA measured. Data are mean (SEM) from 4 animals, and analyzed by ANOVA and Bonferroni post hoc test; values differ from 2-2 within challenge group, * P < 0.05, *** P < 0.001

As supplying glutamine prior to pyolysin challenge supported cytoprotection, we wondered whether glutamine could also help cells recover after pyolysin challenge. Cells were cultured for 24 h in the presence or absence of 2 mM glutamine and then challenged for 2 h with pyolysin, after which the media were replenished with or without 2 mM glutamine for a further 24 h. Cellular DNA was measured at the end of the experiment to estimate cell survival. Cells treated with glutamine prior to pyolysin challenge showed no significant difference (P = 0.18) in cellular DNA remaining when media were replenished with glutamine or not after pyolysin challenge (Fig 3C, 2-2 and 2-0), with cell survival reduced by 26% and 33%, respectively. However, deprivation of glutamine prior to pyolysin challenge, irrespective of whether media were replenished with glutamine or not after pyolysin challenge, reduced cell survival by 70% and 88% (Fig 3C, 0-2 and 0-0, P < 0.001). These data provide evidence that glutamine supported cytoprotection against pyolysin, but glutamine did not help recovery after pyolysin challenge.

### Glutamine supports stromal cell protection against streptolysin O

We next examined whether glutamine affected stromal cytoprotection against another cholesterol-dependent cytolysin, streptolysin O (SLO). We first determined that a 2 h challenge with SLO caused cell damage to bovine endometrial stromal cells, as determined by reduced cell viability and leakage of LDH (Fig 4A, B). However, limiting the availability of glutamine increased the leakage of LDH into cell supernatants when cells were challenged with SLO (Fig 4C; two-way ANOVA; n = 4 animals, P < 0.001). Together the data from Figs 2 to 4 provide evidence that glutamine supports stromal cell protection against cholesterol-dependent cytolysins.

**Fig 4.**
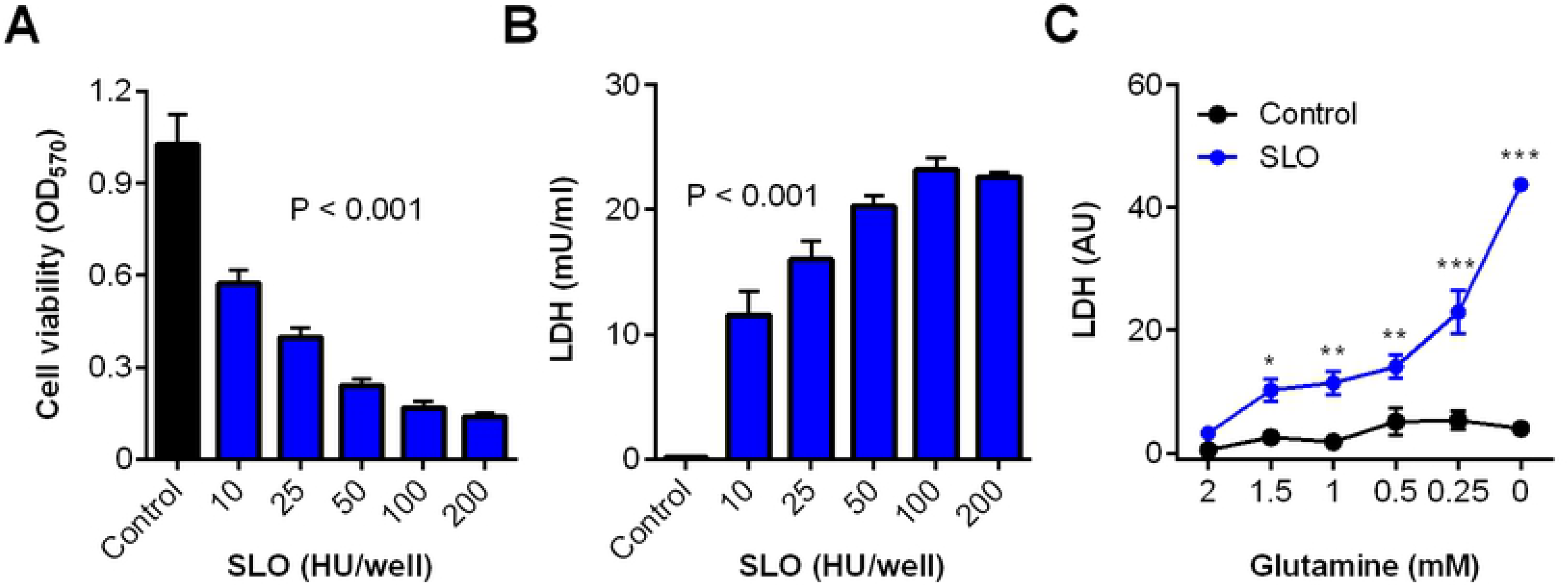
Glutamine is cytoprotection against streptolysin O. Bovine endometrial stromal cells were challenged for 2 h with control serum-free medium (black bar) or medium containing the indicated concentrations of SLO (blue bars); cell viability was determined by MTT assay (A) and LDH leakage measured in the cell supernatants (B). Data are presented as mean (SEM) using cells from 4 animals; data were analyzed by ANOVA and P values reported. (C) Cells were cultured in medium containing the indicated concentrations of glutamine for 24 h, and challenged for 2 h with control medium (•) or 10 HU SLO 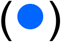, and LDH measured in supernatants and normalized to cellular DNA in the control challenge. Data are mean (SEM) of 4 animals, analyzed by ANOVA with Dunnett’s post hoc test; values differ from 2 mM glutamine SLO challenge, * P < 0.05, ** P < 0.01, *** P <0.001

### Glutamine supports HeLa cell protection against pyolysin and streptolysin O

To examine whether the effect of glutamine on cytoprotection against cholesterol-dependent cytolysins was restricted to bovine endometrial stromal cells, we used immortalized human cervical epithelial cells, HeLa cells, because they are widely employed to examine tissue cell responses to cholesterol-dependent cytolysins [6, 29, 30]. First, we established that challenging HeLa cells with pyolysin caused pore-formation, as determined by a reduction in intracellular potassium after 5 min (Fig 5A), a reduction in cell viability after 2 h (Fig 5B), and an increase in the leakage of LDH from the cytosol into cell supernatants (Fig 5C).

**Fig 5.**
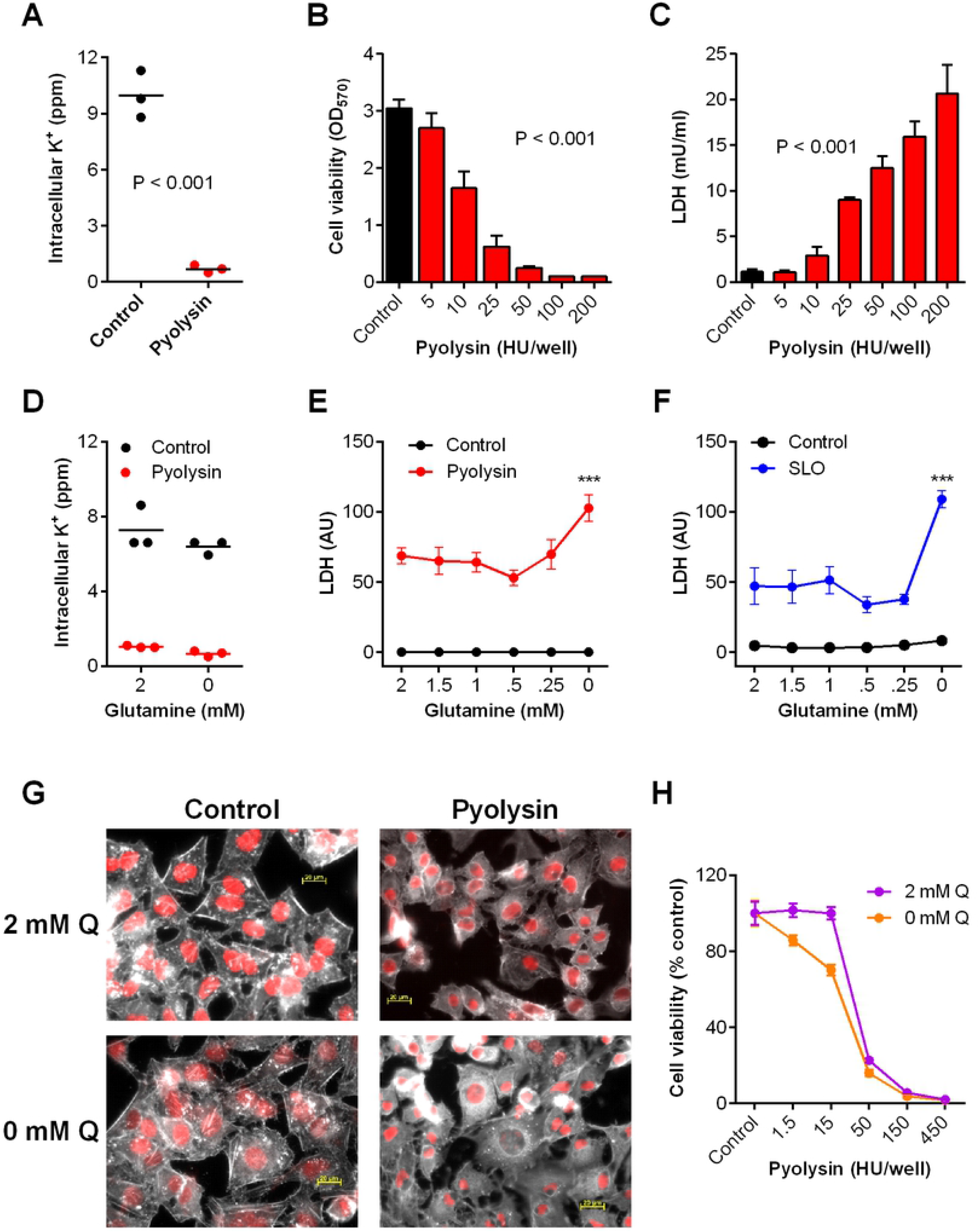
Cytolytic activity of pyolysin in HeLa cells and glutamine. (A) HeLa cells were challenged for 5 min with control serum-free medium (•) or medium containing pyolysin 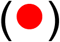, and potassium was measured in cell lysates. Data are presented using cells from 3 independent cell passages and the horizontal line represents the mean; data were analyzed by t-test. (B, C) Cells were challenged for 2 h with control serum-free medium (black bar) or medium containing the indicated concentrations of pyolysin (red bars); cell viability was determined by MTT assay and LDH leakage measured in the cell supernatants. Data are presented as mean (SEM) using cells from 4 passages; data were analyzed by ANOVA and P values reported. (D) Cells were cultured for 24 h in medium containing 2 mM glutamine (2) or without glutamine (0), and challenged for 5 min with control medium (•) or pyolysin 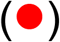. Intracellular potassium was determined by flame photometry. Data are from 3 passages, with the horizontal line indicating the mean. (E, F) Cells were cultured in medium containing the indicated concentrations of glutamine for 24 h, and challenged for 2 h with control medium (•), 10 HU pyolysin 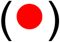 or 10 HU SLO 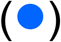, and LDH measured in supernatants and normalized to cellular DNA in the control challenge. Data are mean (SEM) of 4 passages, analyzed by ANOVA with Dunnett’s post hoc test; values differ from 2 mM glutamine cytolysin challenge, *** P <0.001. (G) Cells were cultured for 24 h in the presence of 2 mM glutamine (2 mM Q) or without glutamine (0 mM Q) in serum-free media, and then challenged for 2 h with control medium or pyolysin. The cells were stained with phalloidin to visualize F-actin (white) and fluorescent microscope images collected; nuclei are red; scale bars are 20 μm. Images are representative of 3 experiments. (H) Cells were treated with medium containing 10% FBS with glutamine (2 mM Q, 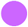) or without glutamine (0 mM Q, 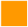) for 24 h before challenge with the indicated concentrations of pyolysin. Cell viability was determined by MTT assay and expressed as the percent of control. Data are mean (SEM) of 4 passages.

To examine if the availability of glutamine affected cytoprotection against pyolysin, HeLa cells were cultured for 24 h in serum-free media containing excess glucose (25 mM) with a range of concentrations of glutamine (0 to 2 mM), and then challenged for 2 h with control medium or 10 HU/well pyolysin. Irrespective of glutamine availability, pyolysin caused pore formation, as determined by loss of intracellular potassium within 5 min (Fig 5D; two-way ANOVA, P < 0.001). However, limiting the availability of glutamine increased the accumulation of LDH in supernatants when HeLa cells were challenge with pyolysin (Fig 5E; two-way ANOVA, P = 0.002). HeLa cells are also sensitive to SLO [30, 31], and we found that glutamine deprivation also increased LDH leakage when HeLa cells were challenges with SLO (Fig 5F; two-way ANOVA, P < 0.0001).

Staining actin with phalloidin also showed that HeLa cells cultured in glutamine lost their characteristic angular shape and became rounded when challenged with pyolysin, although they usually maintained defined cell boundaries (Fig 5G). However, cells deprived of glutamine and challenged with pyolysin were more misshapen with less clear boundaries (Fig 5G).

We also took advantage of similar growth curves for HeLa cells irrespective of the glutamine supply when cells were cultured with 10% fetal bovine serum, as determined by MTT assay (Supplementary Fig 1). Cells cultured with serum but without glutamine prior to pyolysin challenge were more sensitive to cytolysis than cells cultured in 2 mM glutamine (Fig 5H; two-way ANOVA, P = 0.001). Together, the data in Fig 5 provide evidence that glutamine supports HeLa cell protection against cholesterol-dependent cytolysins.

### Glutaminolysis was not essential for cytoprotection against pyolysin

One obvious mechanism for the cytoprotective effect of glutamine against cholesterol-dependent cytolysins is that glutamine could supply cellular energy - even though the cells were supplied with excess glucose (11 mM for stroma, 25 mM for HeLa cells). First we showed that glucose was used by the cells for energy because inhibiting glycolysis with 2-deoxy-D-glucose (2DG) markedly increased LDH leakage from stromal cells challenged with pyolysin, even when cells were supplied with 2 mM glutamine (Fig 6A). To explore the importance of glutamine as an energy substrate for cytoprotection, we examined the role of glutaminolysis, whereby glutaminase converts glutamine to glutamate, which is metabolized to succinate to replenish the Krebs cycle [9, 24]. To inhibit glutaminolysis we used the inhibitor BPTES, a bis-thiadiazole that induces an inactive conformation of glutaminase, and DON, a non-standard amino acid 6-Diazo-5-oxo-L-norleucine that covalently binds glutaminase. We postulated that inhibiting glutaminase in cells supplied with 2 mM glutamine would mimic glutamine deprivation, leading to increased leakage of LDH. As before, in the absence of the glutaminase inhibitors, glutamine deprivation increased the leakage of LDH from cells challenged with pyolysin (Fig 6B, C). However, the leakage of LDH after pyolysin challenge was not significantly increased in cells treated with BPTES (Fig 6B; two-way ANOVA, P = 0.86) or DON (Fig 6C; two-way ANOVA, P = 0.52). In a complementary approach, we cultured cells with a range of concentrations of succinate, in glutamine-free medium, to assess whether the beneficial effect of glutamine was by replenishing the Krebs cycle, and to account for the GABA (gamma-aminobutyric acid) shunt converting glutamine to succinate [9]. However, supplying succinate did not significantly reduce the leakage of LDH from cells in glutamine-free medium (Fig 6D; two-way ANOVA, P = 0.99).

**Fig 6.**
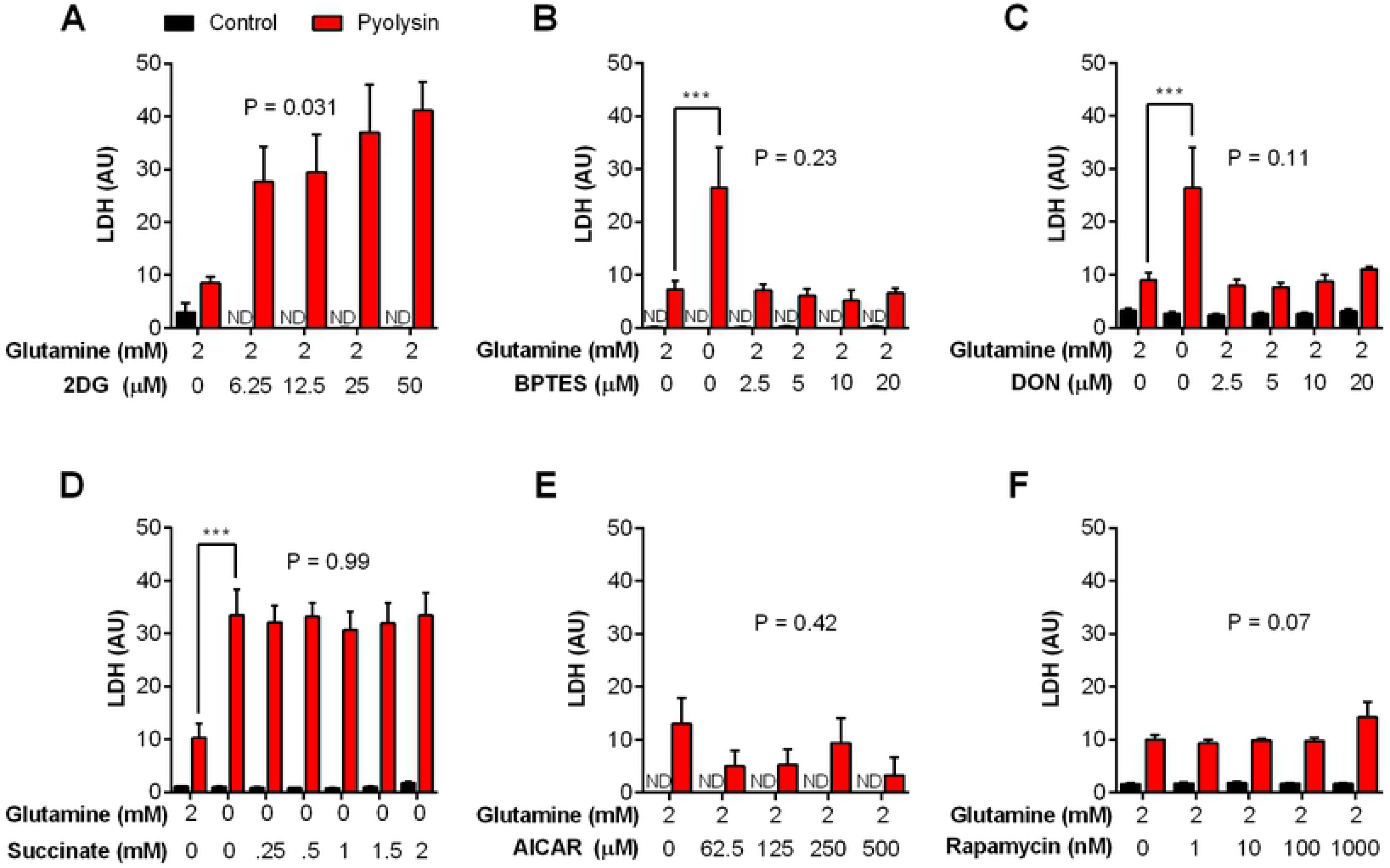
Stromal cell metabolism and protection against pyolysin. Bovine endometrial stromal cells were cultured in serum-free media for 24 h with the indicated concentrations of glutamine and glycolysis inhibitor 2-deoxy-D-glucose (A), glutaminolysis inhibitors BPTES (B) and DON (C), succinate (D), AMPK activator AICAR (E), or mTOR inhibitor rapamycin (F), and then challenged for 2 h with control medium (▀) or pyolysin 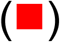. The leakage of LDH from cells was measured in cell supernatants, and normalized to cellular DNA in the control challenge. Data are presented as mean (SEM) using 4 animals for each experiment. Data were analyzed by ANOVA and Dunnett’s post hoc test; reported P values are the effect of the treatment on the response to pyolysin challenge. ND, not detectable.

Cells regulate their energy homeostasis using AMP-activated protein kinase (AMPK), which senses increased AMP:ATP ratios, and mammalian target of rapamycin (mTOR), which integrates satiety signals from hormones, growth factors, and the abundance of amino acids, including glutamine [32, 33]. As glutamine influences AMPK and mTOR signaling [32, 34], we considered whether AMPK and mTOR might affect the ability of glutamine to support cytoprotection against pyolysin. However, there was no substantive effect on LDH leakage from cells challenged with pyolysin when mimicking metabolic energy deficits by activating AMPK with AICAR (Fig 6E; ANOVA, P = 0.42) or inhibiting mTOR with rapamycin (Fig 6F: ANOVA, P = 0.07). Together, the data in Fig 6 provide evidence that cytoprotection against pyolysin was not dependent on glutamine replenishing the Krebs cycle.

### Glutamine and cellular cholesterol

A second mechanism for the cytoprotective effect of glutamine against cholesterol-dependent cytolysins is that glutamine could reduce cellular cholesterol. Methyl-β-cyclodextrin reduces cellular cholesterol [14, 29], and in the present study, treating cells with 0.5 mM methyl-β-cyclodextrin for 24 h reduced cellular cholesterol in HeLa cells (Fig 7A) and in stromal cells (Fig 7B). This reduced cellular cholesterol also protected the cells against a 2 h pyolysin challenge, as determined by MTT assay for HeLa cells (methyl-β-cyclodextrin vs. vehicle, 96.6 ± 7.3 vs 14.8 ± 0.5% viability of control; P < 0.001, t-test, n = 4) and stromal cells (methyl-β-cyclodextrin vs vehicle, 76.7 ± 13.4 vs 11.8 ± 3.2% viability of control; P < 0.001, t-test, n = 7). However, HeLa cell or endometrial stromal cell cholesterol concentrations did not significantly differ when cultured with or without 2 mM glutamine (Fig 7A, B). We also took advantage of the well-defined HeLa cell shape and used filipin and confocal microscopy to examine the distribution of cholesterol [35]. HeLa cells cultured with or without glutamine showed little difference in staining intensity (Fig 7C). Taken together these data do not support the idea that glutamine could alter cytoprotection against cholesterol-dependent cytolysins by modifying cellular cholesterol.

**Fig 7.**
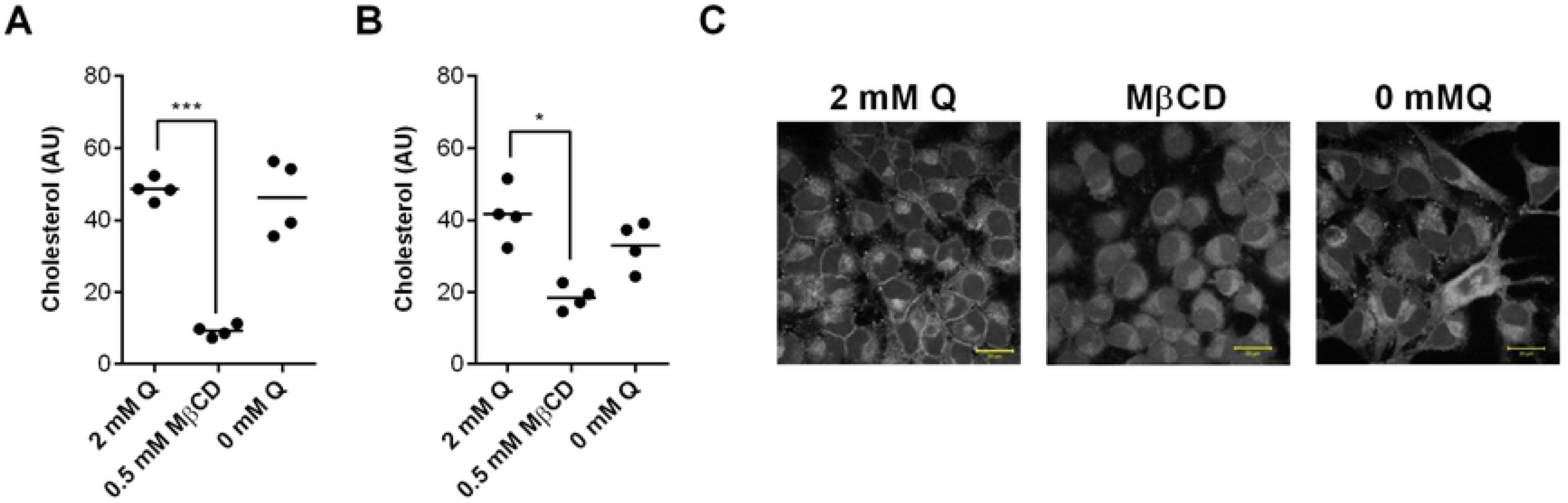
Glutamine and cellular cholesterol. HeLa cells (A) and bovine endometrial stromal cells (B) were cultured for 24 h in serum-free media with glutamine (2 mM Q), 0.5 mM methyl-β-cyclodextrin (MβCD), or without glutamine (0 mM Q). Cellular cholesterol was measured and normalized to the phospholipid content to account for differences in cell growth. Data are presented using 4 independent cell passages for HeLa cells or 4 animals for stromal cells; the horizontal line represents the mean. Data were analyzed by ANOVA and Dunnett’s post hoc test; values differ from 2 mM glutamine, * P < 0.05, *** P < 0.001. (C) Confocal microscope images of HeLa cells stained with filipin to visualize cholesterol (white) for the indicated treatments; scale bars are 20 μm.

## Discussion

We found that glutamine supports the protection of tissue cells against the damage caused by cholesterol-dependent cytolysins from pathogenic bacteria. The role of glutamine in cytoprotection was unexpected, but was consistent across a range of experiments. Glutamine supported cytoprotection against two different cholesterol-dependent cytolysins, pyolysin and streptolysin O, in two disparate tissue cell types, primary bovine endometrial stromal cells and immortalized human cervical epithelial cells.

Even though pores formed in the cell membrane, as evidenced by the rapid leakage of potassium from cells, supplying glutamine reduced the damage that the cytolysins caused to the cells. This glutamine cytoprotection was evident by examining cell viability, imaging cells, and by measuring the leakage of LDH from cells. We chose a 2 h challenge with cholesterol-dependent cytolysins because this would more likely test cytoprotection than a longer cytolysin challenge, which might also reflect longer-term immune and recovery responses [6, 28]. Finding that glutamine reduced pyolysin-induced cell death if given before, but not after pyolysin challenge also supports a role for glutamine in protecting cells, rather than damage repair. The beneficial role of glutamine in tissue cytoprotection complements the role glutamine plays in immune cell metabolism and supporting inflammatory responses to pathogens [9, 36].

Infections are metabolically demanding [9, 25, 37]. We therefore considered whether cytoprotection against cholesterol-dependent cytolysins might depend on glutamine replenishing the Krebs cycle [9]. Although glutaminase is active in fibroblasts and HeLa cells [38, 39], glutaminase inhibitors did not impair cytoprotection against pyolysin in the present study, and supplying succinate to cells in glutamine-free media did not enhance cytoprotection. Damage and infections also activate AMPK, and glutamine regulates AMPK and mTORC1 signaling [32, 34], but manipulating AMPK or mTOR in the present study did not affect cytoprotection against pyolysin. Taken together, these lines of evidence imply that whilst glutamine was important for cytoprotection against cholesterol-dependent cytolysins, the mechanism did not depend on glutaminolysis. These findings are intriguing because although immunity and tolerance complement each other, they can employ different mechanisms [36, 40, 41]. Indeed, whilst succinate did not contribute to cytoprotection against cytolysins here, cellular succinate regulates innate immunity and drives inflammatory responses to bacterial lipopolysaccharide in macrophages [9, 10]. Furthermore, manipulating AMPK or mTOR, as well as reducing the availability of glucose or glutamine, impairs inflammatory responses to lipopolysaccharide in the bovine endometrium [42, 43].

Cholesterol is the binding target for pyolysin and SLO [5], and reducing cholesterol in the cell membrane increases cytoprotection against cholesterol-dependent cytolysins [14, 30, 44]. However, in the present study, any effects of glutamine on cellular cholesterol abundance or distribution were modest compared with methyl-β-cyclodextrin. These findings are plausible because glutamine can provide citrate for cholesterol synthesis, and glutamine stimulates the expression of genes associated with cholesterol synthesis [45, 46].

It was surprising that cells only appeared to need > 0.25 mM glutamine to provide cytoprotection against the cytolysins. Whilst, there is about 0.7 mM glutamine in human plasma and 0.25 mM in bovine plasma [8, 25], it is possible that cells lying within damaged tissues may have less access to glutamine. The need for only a small amount of glutamine may also explain why the glutaminase inhibitors did not increase cellular sensitivity to cytolysins. As glutamine contributes to amino acids, proteins, and nucleotides, as well as supplying the Krebs cycle, future studies might use radiolabeled glutamine to trace where glutamine may contribute to cell protection mechanisms, which include cell stress responses, membrane repair, and cytoskeletal maintenance [6, 7, 15, 28, 29, 31, 47].

In conclusion, we found that glutamine supports the protection of tissue cells against the damage caused by cholesterol-dependent cytolysins from pathogenic bacteria. More work will be needed to determine the mechanism linking glutamine to cytoprotection against cytolysins. However, the implication of finding that glutamine supports cytoprotection is that glutamine may help tissues to tolerate pathogenic bacteria that secrete cholesterol-dependent cytolysins.

## Methods

### Ethical statement

No live animal experiments were performed. Uteri were collected from cattle after slaughter and processing as part of the normal work of a commercial slaughterhouse, with approval (registration number U1268379/ABP/OTHER) from the United Kingdom (UK) Department for Environment, Food and Rural Affairs under the Animal By-products Registration (EC) No. 1069/2009.

### Cell culture

To isolate bovine endometrial stromal cells, uteri were collected after slaughter from post pubertal, non-pregnant animals with no evidence of genital disease or microbial infection. Endometrial stromal cells were isolated, cell purity confirmed, and the absence of immune cell contamination verified, as described previously [14, 48, 49]. Briefly, stromal cells were isolated by enzymatic digestion of the endometrium, sieving the cell suspension through 70-μm mesh to remove debris, and then through a 40-μm mesh (pluriStrainer®, Cambridge Bioscience, Cambridge, UK) to isolate stromal cells, followed by adhesion to culture plates within 18 h, at which time any contaminating epithelial cells were washed away. The cells were maintained in 75 cm^2^ flasks (Greiner Bio-One, Gloucester, UK) with complete medium, comprising RPMI-1640 medium (61870, Thermo Fisher Scientific, Paisley, UK), 10% FBS (Biosera, East Sussex, UK), 50 IU/ml of penicillin, 50 μg/ml of streptomycin and 2.5 μg/ml of amphotericin B (all Sigma).

The HeLa cells (Public Health England; 93021013, HeLa CCL2) were maintained in 75 cm^2^ flasks with complete medium, comprising DMEM (41965, Thermo Fisher Scientific) 10% FBS, 50 IU/ml penicillin, 50 μg/ml streptomycin, and 2.5 μg/ml amphotericin B. The HeLa cell identity was confirmed at the end of the study by short tandem repeat profiling (Report Reference SOJ39361; ATCC, Manassas, VA, USA). Cells were incubated at 37°C in humidified air with 5% CO_2_.

### Cholesterol-dependent cytolysins

The *plo* plasmid (pGS59) was a gift from Dr H Jost (University of Arizona), and pyolysin protein was generated as described previously [14, 50]. The activity of pyolysin was 628,338 HU/mg protein, as determined by hemolysis assay using horse red blood cells (Oxoid, Hampshire, UK), as described previously [14, 51]. Endotoxin contamination was 1.5 EU/mg protein, as determined by a limulus amebocyte lysate assay (LAL endotoxin quantitation kit; Thermo Fisher Scientific, Hertfordshire, UK). Streptolysin O was stored as 1 mg/ml solution and activated using 10 mM dithiothreitol according to the manufacturer’s instructions (Sigma, Gillingham, UK). To examine pyolysin binding, 100 HU/ml pyolysin was incubated for 1 h in PBS with vehicle, 2 mM glutamine, or 1 mM cholesterol, and a hemolysis assay conducted.

### Glutamine manipulation

The bovine endometrial stromal cells were seeded at 5 × 10^4^ cells/well in 24-well plates and incubated for 24 h in complete medium. The cells were then incubated for 24 h in serum-free medium containing the amounts of L-glutamine specified in *Results*, which were generated by combining defined ratios of RPMI1640 with or without L-glutamine (11875, 11.1 mM glucose, 2 mM glutamine; and, 31870, 11.1 mM glucose, no glutamine; Thermo Fisher Scientific).

The HeLa cells were seeded at 4 × 10^4^ cells/well in complete medium for 24 h, followed by a further 24 h in serum-free medium containing the amounts of L-glutamine specified in *Results*, which were generated by combining defined ratios of DMEM with or without L-glutamine (41965, 25 mM glucose, 4 mM glutamine; and, 11960, 25 mM glucose, no glutamine; Thermo Fisher Scientific).

After the treatment period, cells were challenged with their corresponding control medium, or medium containing pyolysin or SLO, as specified in *Results*. In some experiments, transmitted light images of the cells were collected using an an Axiovert 40C inverted microscope and AxioCam ERc5s camera (Zeiss, Jena, Germany). At the end of experiments, cell supernatants were collected for LDH quantification, and cells used for measuring cellular DNA or viability.

### Nutrients and inhibitors

To examine the effect of nutrients, cells were seeded and cultured for 24 h in complete media, and then cultured for 24 h in serum-free media containing the amounts reported in *Results* of dimethyl succinate (W239607, Sigma). After 24 h treatment, cells were challenged for 2 h with pyolysin, and supernatants and cells collected. To examine the effect of inhibitors, cells were seeded and cultured for 24 h in complete media, and then cultured for 24 h in serum-free media containing the amounts reported in *Results* of the glycolysis inhibitor 2-deoxy-D-glucose (2DG; D3179, Sigma), the glutaminase inhibitors BPTES (314045, EMD Millipore, Hertfordshire, UK) or DON (D2141, Sigma), the AMPK activator AICAR (2840, Tocris, Bristol, UK), the mTOR inhibitor rapamycin (553211, EMD Millipore), or methyl-β-cyclodextrin (332615, Sigma). After 24 h treatment, cells were challenged for 2 h with control medium or pyolysin, and supernatants and cells collected.

### Cell viability

The mitochondria-dependent reduction of 3-(4,5-dimethylthiazol-2-yl)-2,5-diphenyltetrazolium bromide (MTT, Sigma) to formazan was used to assess cell viability, as described previously [14]. As nutrient availability may influence the reduction of MTT, cell abundance was also determined by measuring cellular DNA content. Briefly, at the end of experiments when supernatants were removed, the cells were washed in 500 μl ice-cold PBS before being stored at −80°C overnight to ensure lysis, and DNA was measured using the CyQUANT Cell Proliferation Assay Kit (Thermo Fisher Scientific).

### Lactate dehydrogenase and potassium leakage

Lactate dehydrogenase leakage from cells was measured in cell supernatants using a Lactate Dehydrogenase Activity Assay Kit (Cambridge Bioscience) [6, 51]. Where indicated in *Results*, LDH leakage from cells was normalized to the cellular DNA in the control challenge.

To examine potassium leakage, 7.5 × 10^5^ cells were seeded in 75 cm^2^ culture flasks in complete media for 24 h, before treatment with or without 2 mM glutamine for a further 24 h in serum-free media. Media were then discarded and cells washed 3 x with potassium-free choline buffer (129 mM choline-Cl, 0.8 mM MgCl_2_, 1.5 mM CaCl_2_, 5 mM citric acid, 5.6 mM glucose, 10 mM NH_4_Cl, 5 mM H_3_PO_4_, pH 7.4; all Sigma). Cells were then incubated in choline-buffer with control medium or pyolysin for 5 min at 37°C. Subsequently, cells were washed 3 x in ice-cold choline-buffer and lysed in 0.5% Triton X-100 (Sigma) in double-distilled water for 20 min at room temperature with gentle agitation. Potassium was measured in the cleared lysates using a Jenway PFP7 flame photometer (Cole-Parmer, Stone, Staffordshire, UK).

### Cholesterol assay

The bovine endometrial stromal cells and HeLa cells were grown at a density of 10^5^ cells/well in 12-well tissue culture plates for 24 h in complete media, and then cultured for 24 h in serum-free media with or without 2 mM glutamine, or 0.5 mM methyl-β-cyclodextrin, as described in *Results*. After the treatment period, cells were collected in 200 μl/well cholesterol assay buffer (Thermo Fisher Scientific) and stored in Eppendorf tubes at −20°C. When needed, samples were defrosted at room temperature and sonicated for 10 min in a sonicating water bath. Cellular cholesterol content was measured using the Amplex® Red Cholesterol Assay Kit (Thermo Fisher Scientific). Total cellular phospholipid was measured in the samples prepared for the cholesterol assay using a phospholipid assay kit (MAK122, Sigma). Cholesterol concentrations were then normalized to phospholipid concentrations.

### Immunofluorescence

To examine actin distribution, cells were seeded at a 5 × 10^4^ cells on glass coverslips in a 24-well plate in complete medium for 24 h, followed by a further 24 h in serum-free medium with or without 2 mM glutamine. Cells were challenged for 2 h with the corresponding control medium or medium containing pyolysin, as specified in *Results*. Cells were washed with PBS, fixed with 4% paraformaldehyde, washed in PBS and then permeabilized in 0.2% Triton X-100. Cells were then blocked using 0.5% bovine serum albumin and 0.1% Triton X-100 in PBS, followed by incubation with Alexa Fluor™ 555 Phalloidin (Thermo Fisher Scientific). Cells were washed in 0.1% Triton X-100 in PBS three times and mounted onto microscope slides, using 4′,6-diamidino-2-phenylindole (Vectashield with DAPI; Vector Laboratories Inc., Burlington, CA, USA) to visualize cell nuclei. Cell morphology and target localization were analyzed with an Axio Imager M1 upright fluorescence microscope (Zeiss, Jena, Germany) and images captured using an AxioCamMR3.

To image cholesterol, 5 × 10^4^ cells were seeded on glass coverslips in a 24 well plate in complete medium for 24 h, followed by 24 h in serum-free medium with or without L-glutamine, or 0.5 mM methyl-β-cyclodextrin, as described in *Results*. Coverslips were washed with PBS, fixed with 4% paraformaldehyde, and washed with PBS. Coverslips were then incubated for 45 mins at room temperature with 50 μg/ml filipin III from *Streptomyces filipinensis* (Sigma). Cells were washed with PBS before being mounted using 2.5% Mowiol mounting medium containing 2.5% DABCO (1,4-diazabicyclo-(2,2,2)-octane, Merck). Cell cholesterol was analyzed using a LSM710 confocal microscope (Zeiss) with the Zeiss Zen 2010 software. Images were captured using a × 63 oil objective using the channel range 410-476 nm, and coverslips were subjected to identical exposure times and conditions.

### Statistical analysis

Data are presented as arithmetic mean and error bars represent SEM. The statistical unit was each animal used to isolate bovine endometrial stromal cells or each independent passage of HeLa cells. Statistical analysis was performed using SPSS 22.0 (SPSS Inc. Chicago, IL), and P < 0.05 was considered significant. Comparisons were made between treatments using one-way or two-way ANOVA with two-tailed Bonferroni or Dunnett posthoc test, or unpaired two-tailed Student’s t test, as specified in *Results* and figure legends.

## Acknowledgements

We thank J Cronin and E Dudley for advice, T Ormsby for technical assistance, and H Jost for supplying pyolysin.

## Supporting information

**SI Fig 1. Similar cell growth curves irrespective of glutamine supply for HeLa cells cultured with serum**. HeLa cells were cultured in medium containing 10% fetal calf serum and 2 mM glutamine for 24 h, and then with or without 2 mM glutamine for a further 72 h. Cell viability was measured using the MTT assay every 24 h. The data are reported as mean (SEM) from 4 independent passages. Data were analyzed by 2-way ANOVA; there was a significant effect of time (F_(3, 24)_ = 113.6, P < 0.0001) but not for glutamine (F_(1, 24)_ = 0.0005, P = 0.98) or the interaction of time × glutamine (F_(3, 24)_ = 0.5, P = 0.71).

